# Tosylation-mediated uronic acid knockout from cryoprotective FucoPol revealed the importance of polyanionicity in ice growth disruption

**DOI:** 10.1101/2025.08.27.672662

**Authors:** B. M. Guerreiro, J. C. Lima, J. C. Silva, F. Freitas

## Abstract

Negative formal charge is a critical structural trait for the expression of cryoprotection due to its strong ice growth disruption at the growing ice facet. Polyanionic polysaccharides containing uronic acids (UA) in their structure are therefore often associated with psychrophilic-like traits, such as defense against cryoinjury. However, their practical biological post-thaw benefit is convoluted amongst other contributing factors, including hydrophilic-hydrophobic balance, molecular weight, rheological properties, and structural conformation. Here, we leveraged ubiquitous (31.7% m.d.w.) and COOH-targeted (63.4% m.d.w.) leaving-group SN_2_ tosylation procedures, coupled with clean purification methodologies (high-MW membrane dialysis), to synthesize UA-free variants (FP-OTs^18^, FP-OTs^96^) of the cryoprotective, UA-containing polysaccharide FucoPol (FP). Tosylation was confirmed by *ATR*-FTIR (1602.3 cm^−1^ shift to 1627.9 cm^−1^) and ^1^H-NMR (brs, *δ* 7.92 ppm; brs, 7.53 ppm). Tosylated derivatives exhibited reduced solution conductivity (Ω=300 *µ*S/cm vs. ~800 *µ*S/cm), unchanged molecular weight (M_*w*_), and retained biocompatibility. However, the substitution of hydrophilic, low-steric hindrance carboxyl groups with hydrophobic, high steric hindrance tosyl groups promoted a 30–40% reduction in zero-shear viscosity (*η*_0_) and precluded the formation of a hypothermic gel-state due to loss of inter-chain non-covalent interactions, leading to a drastic loss of cryoprotective activity (10–70% of the original post-thaw viability observed for native FucoPol). The cryobiologically antagonistic effect of FP-OTs^18^ and FP-OTs^96^ reveals that UAs are a key player in polysaccharide-based cryopreservation, not only due to standalone beneficial polyanionicity, but also because their presence largely contributes to the expression of agonistic structural traits that elicit a cryoprotective response.

## Introduction

The central dogma of chemical cryobiology is the need for cryoprotectants to possess a net dipole moment in order to exert ice growth disruption by interacting with stable hydrogen bonds necessary for crystal periodicity [1]. A recent multidimensional meta-analysis of perceived structure-function relationships in extremophilic polysaccharides revealed that net formal charge, particularly the polyanionicity conferred by uronic acids (UA), is a critical structural trait towards the expression of several properties, such as antioxidant, bioflocculant and cryoprotective effects [2]. This electrostatic interplay becomes especially relevant in mechanisms of cryoinjury avoidance, such as efficient binding at the liquid-solid interface of ice. The electrostatic disruption of ice growth by negatively charged species, whether permanently or by transient dipoles, is a core aspect in the expression of cryoprotective traits [3, 4]. In small molecular cryoprotectants such as glycerol and DMSO, the role of anionicity is drastically evident [5, 6], but a compounded effect of net charge, hydrophilic-hydrophobic balance, diffusional kinetic hampering induced by molecular weight and varied molecular conformations, among others, in polysaccharides, clouds this evident contribution. A recent study which screened several polysaccharides as potential candidates for cryopreservation formulas demonstrated that polyanionicity was essential but not the sole contributive factor towards the expression of increased post-thaw survival [7]. To further understand the impact of an absent polyanionic character in cryoprotective function, we leveraged our understanding of the fully elucidated chemical structure of FucoPol, a bio-based fucose-rich polysaccharide produced by the Gram-negative bacterium *Enterobacter* A47 (DSM 23139) [8], which has been the subject of extensive cryoprotective research [7, 9–13], to perform a structural UA knockout using organic chemistry methodologies. FucoPol is a high-molecular-weight (1.75.8×10^6^ Da) fucose-containing cryoprotective extracellular polysaccharide (EPS) with a fucose, galactose, glucose, and glucuronic acid hexamer motif (2.0:1.9:0.9:0.5 M ratio), a main chain composed of a →4)-*α*-L-Fuc*p*-(1→4)-*α*-L-Fuc*p*-(1→3)-*β*-D-Glc*p*(1→ trimer repeating unit, and a trimer branch *α*-D–4,6-pyruvyl-Gal*p*-(1→4)-*β*-D-GlcA*p*-(1→3)-*α*-D-Gal*p*(1→ in the C_3_ of the first fucose [9]. It also contains 13–14 wt.% pyruvyl, 3– 5 wt.% acetyl, and 2–3 wt.% succinyl in its composition [8]. The presence of glucuronic acid as well as the acyl substituents pyruvyl and succinyl, confer a polyanionic character to the biopolymer [8]. FucoPol has so far demonstrated strong non-colligative ice growth disruption and re-shaping [9], nucleation promotion and stochastic narrowing [13], versatile use in cryoprotective formulas of variable composition and different cell lines [10], and an antioxidant effect [14, 15] that counteracts the reactive oxygen species-mediated, cryopreservation-induced MPTP opening that leads to delayed post-thaw cell death [16]. All these properties, associated with a high molecular weight, high viscosity and shear-thinning behavior [17] that facilitates nutrient diffusion and promotes cellular attachment by natural design [18], conceive a molecule capable of maximizing the post-thaw viability of cryopreserved biologicals. Therefore, we have conducted the removal of intrinsic, structural UA moieties from native FucoPol (FP) using different leaving-group SN_2_ tosylation procedures. The biological properties of the tosylated products (FP-OTs) were then re-assessed to ascertain the influence in its previously proven cryoprotective performance. This complex structure-function effect convolution in polymeric molecules has remained the greatest challenge in dissecting a singular contributor to the overall expression of a biological effect, because *e*.*g*. the removal of a structural constituent may impact other properties in unperceived ways which ultimately may affect its final function. For this reason, other polysaccharide functionalizations across the literature retain a high degree of simplicity, such as a simple removal of phosphate groups from the structure to assess function [19]. Ultimately, this analysis remains necessary towards a deeper understanding of polysaccharide cryoprotection, and polysaccharide functionalization provides a unique approach to studying the consequences of small structural variations in antifreeze activity. An efficient tosylation of FucoPol also creates an opportunity to generate a building-block structure to anchor other bioactive molecules onto the polymer surface because the tosyl moiety is an excellent leaving group.

## Materials & Methods

### General procedures for the organic synthesis of tosylated FucoPol (FP-OTs)

#### Ubiquitous non-targeted tosylation

In a round-bottom flask, a solution of native FucoPol 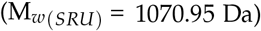 was dissolved in water (volume to make a 4.67 M solution of substrate) and homogenized for 1 h under magnetic stirring. Then, tosyl chloride (M_*w*_ = 190.65 Da, 7 equiv.) and triethylamine (M_*w*_ = 101.193 Da, 7 equiv.) were added, and the reaction mixture was stirred, first at 4 °C for 10 min, then at r.t. for 21 h in a N_2_(g) atmosphere. After complete consumption of the starting material, the reaction mixture was separated into two equivolumetric flasks, designated fraction *1* and fraction *2*. Fraction *1* was concentrated under vacuum, washed three times with ethanol and dried. Fraction *2* was concentrated under vacuum, attempted to precipitate with 0.5 volumes of THF unsuccessfully, dried, washed three times with ethanol, centrifuged at 2,830 *× g* for 5 min and dried again. Fibers resembling native FucoPol were denoted with a suffix *a*, and spherical aggregates with the suffix *b*.

##### Compound 1a

Following the general procedure, from 250 mg of FucoPol, compound *1a* was obtained in 23.1% yield 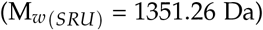; Appearance changed from beige dense cotton fiber to a slightly more transparent beige thin powder-ish film. IV (ATR) *ν*_max_ (cm^−1^) 1655, 1628. ^1^H NMR (500 MHz, D_2_O) *δ* 7.92 (brs, 1H), 7.70 (s, 2H), 7.53 (brs, 1H), 7.38 (s, 2H).

##### Compound 1b

Following the general procedure, from 250 mg of FucoPol, compound *1b* was obtained as part of a 23.1% yield of compound *1a*; Solid gel-like yellow aggregates thought to be TEA and p-toluenesulfate salts enclosed by FP-OTs fibers. IV (ATR) *ν*_max_ (cm^−1^) 2985, 2737, 2702, 1655, 1628, 1475. ^1^H NMR (500 MHz, D_2_O) *δ* 7.92 (brs, 1H), 7.70 (s, 2H), 7.53 (brs, 1H), 7.38 (s, 2H).

##### Compound 2b

Following the general procedure, from 250 mg of FucoPol, compound *2b* was obtained in 8.1% yield; Solid gel-like yellow aggregates thought to be TEA and p-toluenesulfate salt precipitates. IV (ATR) *ν*_max_ (cm^−1^) 2985, 2737, 2702, 2620, 2605, 2531, 2498, 1655, 1628, 1475. ^1^H NMR (500 MHz, D_2_O) *δ* 7.70 (d, *J* = 7.8 Hz, 1H), 7.38 (d, *J* = 7.9 Hz, 1H).

#### C_6_–COOH targeted derivatization

To target the substitution of the carboxyl –OH group, a 1:1:1:1 molar ratio of COOH:FP_*SRU*_ was used. To a 0.25 wt.% native FucoPol solution (500 mg, 1.33*×*10^−4^ mmol) containing 0.467 mmol of –COOH, 0.467 mmol of TsCl (89 mg) and TEA (65.08 *µ*L) were added. To investigate varying degrees of substitution, the reaction was left stirring at room temperature for 18 and 96 h. Then, the end product was purified by dialysis, using cellulose semipermeable membrane strips with distilled water as counter-exchange solvent. The extent of purification was monitored by measuring the pH and conductivity (Ω) both inside and outside the dialysis strips over several rounds of solvent exchange and considered complete when a plateau in both parameters was reached. The final solution was frozen to –80 °C, freeze-dried and dry-weighed to obtain reaction yield.

#### *ATR*-FTIR

Attenuated Total Reflectance Fourier-Transform Infrared Spectroscopy (*ATR*-FTIR) data was recorded on a PerkinElmer Spectrum Two (Waltham, MA, USA) in the range of 4000–400 cm^−1^, by compressing ca. 3–5 mg of freeze-dried polymer with the instrument lever.

#### ^1^H-NMR

Proton Nuclear Magnetic Resonance (^1^H-NMR) spectra were recorded on a Bruker ARX400 (Billerica, MA, USA) at 500 MHz. Chemical shifts are reported in parts per million (ppm, *δ* units). The following abbreviations were used to describe the spectral peaks: s: singlet, d: doublet, t: triplet, q: quartet, m: multiplet, brs: broad singlet.

#### Rheology

Rheological analysis was performed on a MCR92 modular compact rheometer (Anton Paar, Graz, Austria) equipped with a cone-plate geometry (angle 2°, diameter 35 mm, 0.145 mm gap). Temperature was controlled by an in-built Peltier system and set to 37, 20, or 4 °C.

##### Apparent viscosity

Each polysaccharide was dissolved in deionized water, each solution being magnetically stirred for at least 2 h at ambient temperature until homogeneous dissolution was achieved. Flow curves were determined using a steady-state flow ramp in the shear rate range of 10^2^–10^4^ s^−1^. Steady-state tests were carried out for polysaccharide solutions of three different concentrations. To determine the zero-shear viscosity *η*_0_, the Cross model [20] was used to fit the flow curve data:

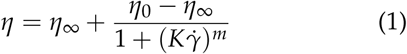

where *η* is the apparent viscosity (mPa·s), *η*_∞_ is the apparent viscosity of the second Newtonian plateau (mPa·s), here considered negligible (*η*_∞_ ≪ *η*_0_ and *η*), *η*_0_ is the zero-shear rate viscosity, 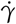 is the shear rate (s^−1^), *K* is a time constant (s) and *m* is the Cross exponent.

##### Oscillatory shear tests

Amplitude sweep experiments (0.1–100%, 1 Hz) were performed to determine the viscoelastic region (LVE) of the polysaccharide solutions. Frequency sweep tests were carried out between 1 and 100 Hz using a predetermined constant strain (0.5–5%) for each sample within the LVE region. Storage (*G*′) and loss (*G*″) moduli were determined for polysaccharide solutions under different concentrations. The gel point was determined at the intersection between *G*′ and *G*″ curves. Gel strength (GS) was determined as the area under the curve delimited by the *G*′ and *G*″ curves, after gel point crossover:

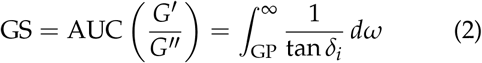

where GP is the frequency at which a gel point was observed and *δ*_*i*_ corresponds to the loss factor at every frequency *i*.

#### *In vitro* methodology

Vero cells (monkey kidney epithelium, ATCC^®^ CCL–81™) were used as the standard cell line, according to ISO 10993–5:2009, for the determination of polysaccharide cytotoxicity and cryoprotective activity.

##### Cell culture

Vero cells frozen in a cryovial containing 10% glycerol and 10% FBS dissolved in a phosphate-buffered saline solution (PBS) were thawed in a water bath at 37 °C for a maximum elapsed time of 2 minutes. Then, cells were centrifuged at 100*×g* for 10 minutes. The supernatant was discarded, cells were resuspended in low-glucose DMEM and transferred to a T-flask. Cells were maintained in incubation at 37 °C in a 5% CO_2_ atmosphere. Cell experiments were carried out when Vero cells achieved 70–80% cell confluency.

##### Cytotoxicity assay

Vero cells were seeded in 96-well microplates containing 100 *µ*L DMEM, at a concentration of 10,000 cells/cm^2^ and left to incubate for 24 h at 37 °C in a 5% CO_2_ atmosphere. After cell adherence, the culture medium was discarded, cells were washed with PBS (Ca^2+^, Mg^2+^–free) and exposed to all test polysaccharides, a positive control (DMEM) and a negative control (100% DMSO). Again, they were left to incubate under the same conditions. After 48 h, well volumes were discarded and 100 *µ*L of 0.08 mg/mL resazurin dissolved in DMEM was added and incubated for 2 h for metabolic viability determination.

##### Cell monolayer cryopreservation

Vero cells were seeded in 96-well microplates with a cell density of 40,000 cells/well in low-glucose DMEM, and incubated at 37 °C, 5% CO_2_ for 24 h to achieve 70–80% cell confluency before freezing. Then, the adhered cells were exposed to a freezing medium containing: 10% (v/v) DMSO and each test polysaccharide at a given concentration, dissolved in DMEM, for a total volume of 100 *µ*L in each well. Polysaccharide concentration gradients were prepared by two-fold serial dilution, ranging from 1.0–0.125 wt.%. The addition of DMSO was performed in the solution preparation phase rather than direct microwell injection to avoid in-well DMSO heterogeneity and minimize pre-freeze exposure time. After medium exposure, the microplate was covered in parafilm and frozen at –80 °C in a Styrofoam container designed to yield an optimal cooling rate of –1 °C/min (*±*0.2 °C). After 24 h, the microplate was thawed in a 37 °C water bath for 2 min. Each well content was diluted 3-fold with PBS and left to incubate at 37 °C, 5% CO_2_ for another 24 h, for cells to regain normal metabolic rate and to avoid an overestimation of viability due to post-thaw cellular apoptosis events that occur around the 6-hour mark post-thaw [21]. The cryoprotective performance of each polysaccharide is a relative metric, contrasting the performance of 10% (v/v) DMSO. The strategic co-supplementation of 10% (v/v) DMSO was designed to compensate for the shortcomings of non-cryoprotective polysaccharides and retain considerable cell recovery rates for statistical significance testing.

##### Post-thaw metabolic viability

After the thawing procedure described for both cytotoxicity and cryopreservation experiments, the liquid content of all wells was discarded and 100 *µ*L of the metabolic indicator resazurin (0.08 mg/mL) were added. The microplate was left to incubate at 37 °C, 5% CO_2_ for 2 h. Absorbance was measured at 570 and 600 nm in a BioTek^®^ microplate reader and visualized in the BioTek^®^ Gen 5 software. Resazurin in the absence of cells was used as experimental control. The commercial cryogenic formulation CryoStor™ CS5 and 10% (v/v) DMSO were used as positive control. DMEM was used as negative control.

##### Post-thaw cell histology

After resazurin exposure and data collection, the liquid contents of all wells were discarded. Image acquisition of post-thaw cell histology was performed with a NIKON Eclipse Ti-S optical microscope connected to a NIKON D610 digital camera, using the Camera Control Pro software.

## Results

Figure 1 summarizes the two synthesis pathways developed for the efficient tosylation of the cryoprotective polysaccharide FucoPol and subsequent chemical and biological analysis. Ultimately, three different contrasting products were obtained for biological cell cryopreservation: native FucoPol (FP) and two tosylated derivatives, FP-OTs^18^ and FP-OTs^96^.

**Figure 1:**
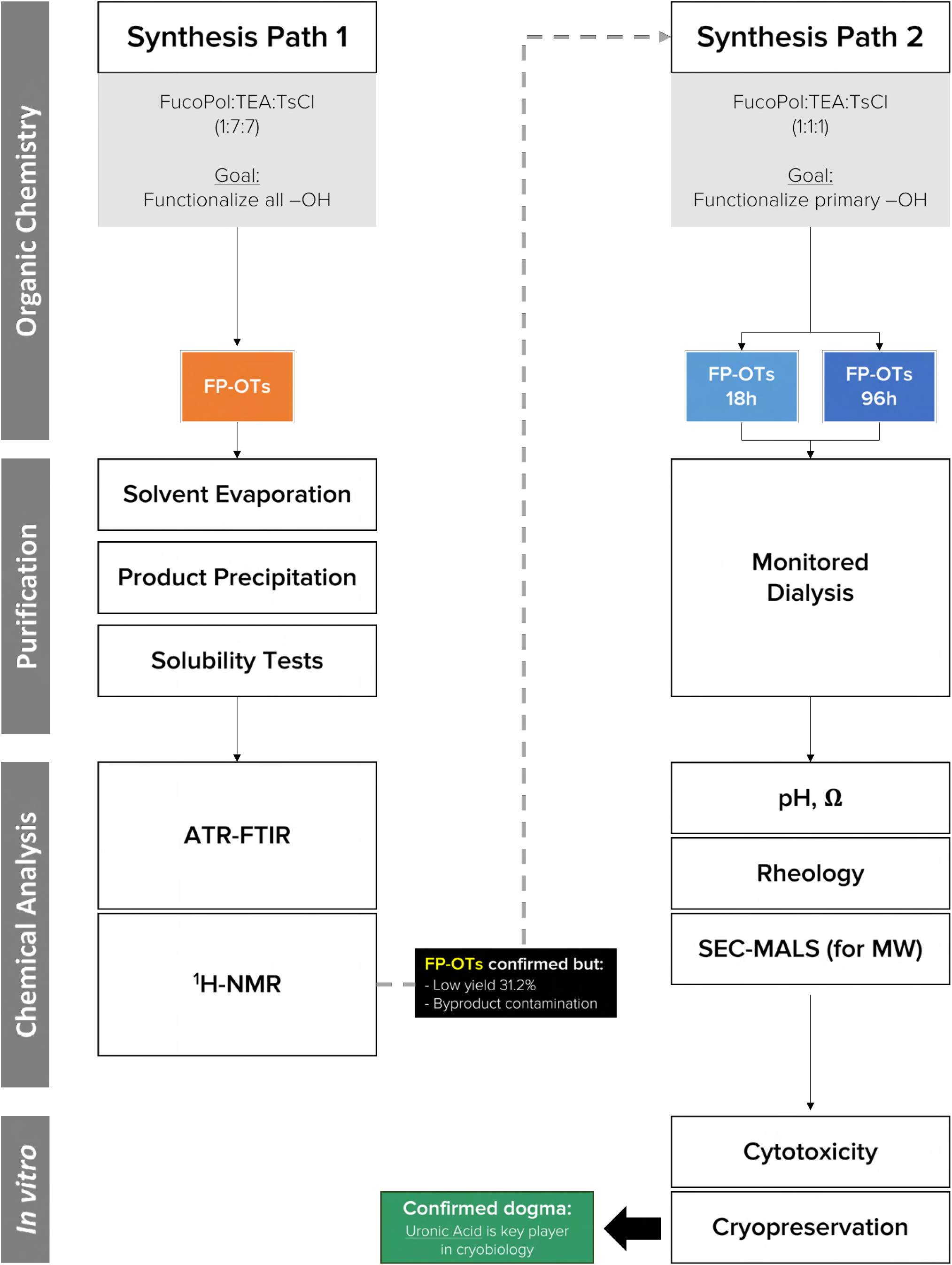
Rationale for the uronic acid knock-out organic synthesis and biological validation experiment.

### FP-OTs synthesis pathway 1: ubiquitous tosylation

The first attempt at FucoPol derivatization was carried out by implementing excess molar equivalents of tosyl chloride (TsCl) and triethylamine (TEA) to establish proof-of-concept verification that tosylation was possible in polysaccharide solutions prone to chain entanglement and viscosity increase which would reduce diffusion-dependent chemical reactions. Therefore, a 1:7:7 ratio of the molar amount of structural repeating unit of FucoPol (FP_*SRU*_), TsCl and TEA was mixed under stirring at 4 °C for the first 10 minutes, then at room temperature for 21 h. Figure 2 summarizes the reactional scheme for both synthetic pathways performed in this study. The ubiquitous tosylation procedure (pathway 1) is a non-specific derivatization attempt of all –OH groups, labeled with **1** on the diagram, regardless of reactivity. Due to solution viscosity possibly playing a role in reactivity, a conservatively diluted 0.25 wt.% aqueous solution of native FucoPol was used. Figure 3 shows the macroscopic evolution of the reaction mixture over time. The initial solution pH was 6 (Figure 3a). The beige turbid color of a homogeneous polymer solution immediately changed to a yellow tint upon addition of TEA (Figure 3b). After 2 h, the reaction mixture was heterogeneous, some condensation was observed, turbidity was less apparent and some insoluble particulate matter suspected to be unreacted TsCl was visible, which was corroborated by a pH 12 measurement (Figure 3c). After 21 h, the reaction was considered to be complete: the characteristic turbidity of FucoPol is present although less intense (Figure 3d) and some precipitate is clearly visible, believed to be excess TsCl and TEA in the form of insoluble triethylammonium p-toluenesulfonate salts. The final reaction mixture had pH 4, believed to be due to excess p-toluenesulfonic acid and HCl byproducts.

**Figure 2:**
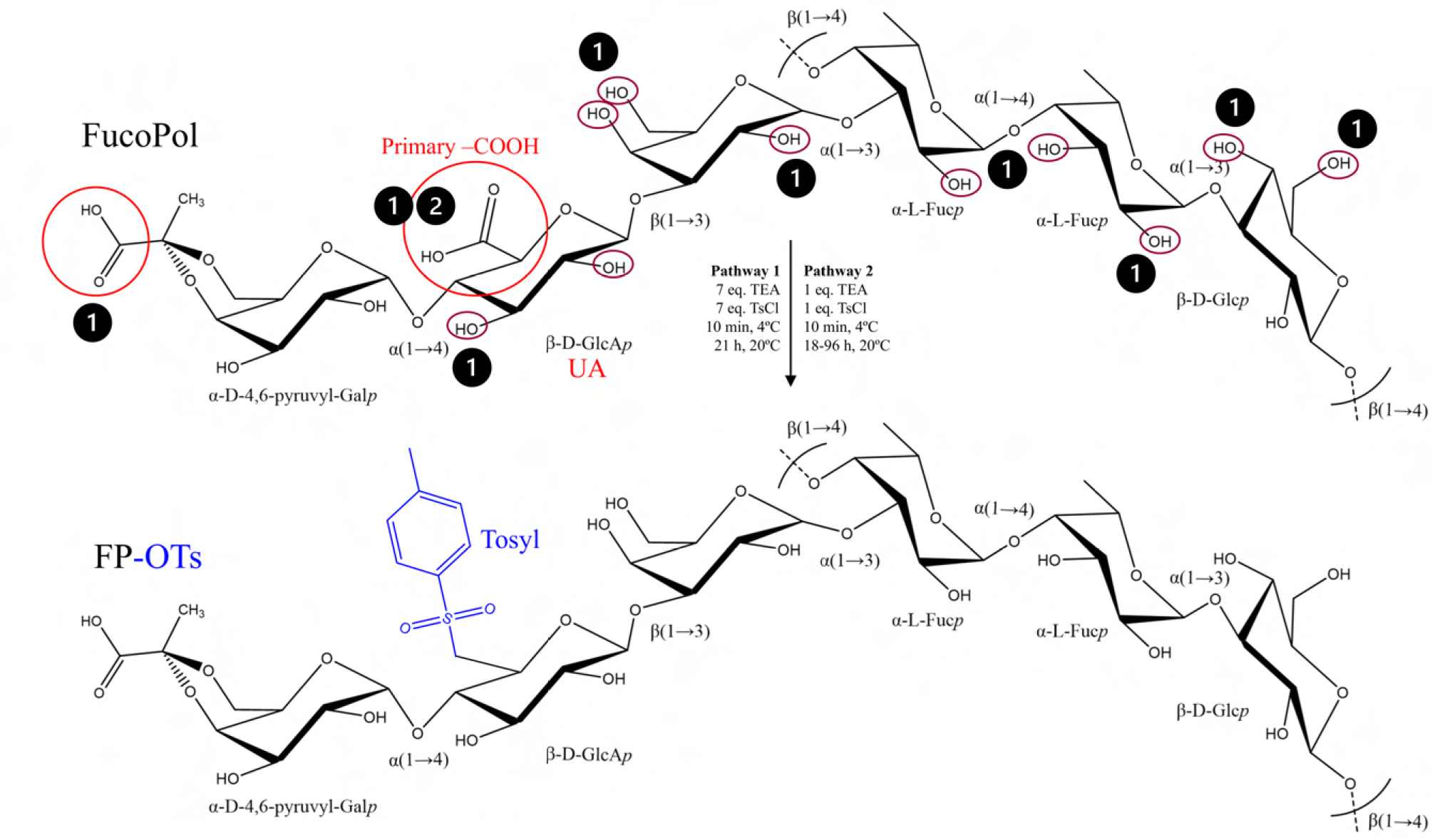
Reactional scheme for the tosylation of FucoPol, by ubiquitous (**1**) and C6-COOH-targeted (**2**) tosylation. The red bubble denotes the only primary carboxyl group of glucuronic acid (GlcA), the only uronic acid, in the structural repeating unit of FucoPol (FP_*SRU*_). The intended product, tosylated FucoPol (FP-OTs), is shown below, with the tosyl leaving group highlighted in blue.

**Figure 3:**
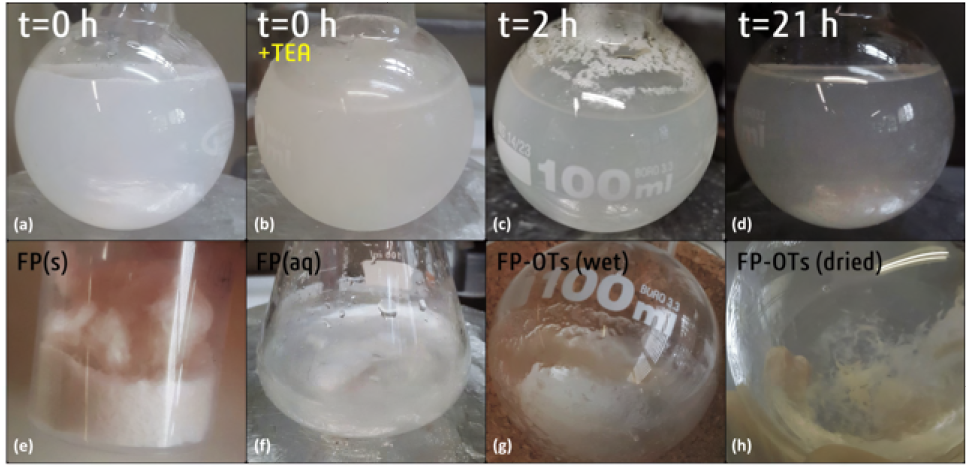
Temporal evolution of the appearance of a 0.25 wt.% FucoPol solution during ubiquitous tosylation (pathway 1). The beige opaque color of an aqueous FucoPol solution in the presence of TsCl (a), otherwise translucent (f), immediately changed to yellowish upon addition of triethylamine (b) and lost turbidity after 2 h (c). After 21 h, any excess TsCl presented as a precipitate of triethylammonium p-toluenesulfate (d). The appearance of each solid form is shown in the bottom panel: native FucoPol fibers (e), FP-OTs post ethanol washing (f) and solvent rotary evaporation (g), FP-OTs post vacuum drying (h), revealing two sub-products: transparent film fibers (compound *1a*) and solid yellow aggregates (compound *1b*, visually similar to compound *2b*).

#### Solvent evaporation

The final synthesis product (Figure 3d) was split into two liquid fractions to assess different purification strategies. Both fractions were initially concentrated under vacuum using a rotary evaporator (Figure 4) to minimize processing volume for further purification. Both fractions resulted in a beige, semi-translucent liquid that slowly transformed into a slightly viscous, opaque, milky substance just before changing into a gel-like substance centrifugally adhered to the walls of the flask (Figure 3g).

**Figure 4:**
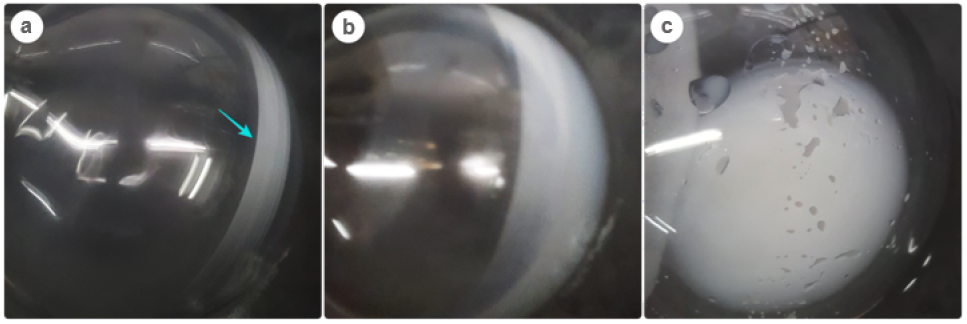
Stages of a tosylated FucoPol solution in a rotary evaporator. The liquid-like solution (a) becomes viscous and milky (b) with increased solvent evaporated (c).

#### Organic solvent washing

After evaporation, different strategies for organic solvent washing were employed for each fraction. Fraction *1* was washed three times with ethanol and dried, resulting in a 21.3% yield. The dry product appearance (Figure 3h) changed from cotton-like fibers (Figure 3e) to smaller strands of the same color (compound *1a*), although a different morphology may be due to only native FucoPol having been freeze-dried. A yellowish solid aggregate was also observed, believed to be precipitated byproduct salts adsorbed onto polymer fibers (compound *1b*). Fraction *2* did not evaporate efficiently and was unsuccessfully precipitated by washing with 0.5 volumes of THF (forming a two-phase system, with a murky dark yellow substance layered on top and surrounded by the solvent) or triple washing with ethanol alone. Precipitation was successful by vigorous agitation in a 1:1 THF:H_2_O mixture, followed by triple washing with ethanol and centrifuging for 5 min at 2,830*×g*, from which the supernatant was discarded, only the pellet being collected and vacuum dried. The product was also a yellowish aggregate (compound *2b*). Ultimately, organic solvent evaporation and washing techniques resulted in a total 31.2% yield for ubiquitous tosylation.

#### Product precipitation

Additional strategies for the one-step purification of FP-OTs were tested. Figure 5 shows the results of Buchner liquid-solid filtration followed by TLC silica plate chromatography for confirmation. A TLC silica plate spiked with the eluent carrying FP-OTs showed no signs of sugar blotting after revealing with a mixture of 1:1 H_2_SO_4_:methanol (Figure 5a), which indicated that FP-OTs did not percolate the filter. In fact, FP-OTs cannot be filtrated by Buchner extraction because the polymer fibers adhere to the cellulose filter and become practically indissociable, as observed by a white wax-like texture on top of the cellulose filter (Figure 5b). Contrary to most substances, which form easily scrapable crystals, polymeric fibers become compacted against the cellulose layer under the action of vacuum. Subsequent pressurized liquid elution becomes completely hampered, thus Buchner liquid-solid extraction is unfeasible. After applying the same revealing procedure, most of the cellulose reacts and becomes charred, but some brown spots appear, indicating the localized sugar decomposition of FP-OTs (Figure 5c).

**Figure 5:**
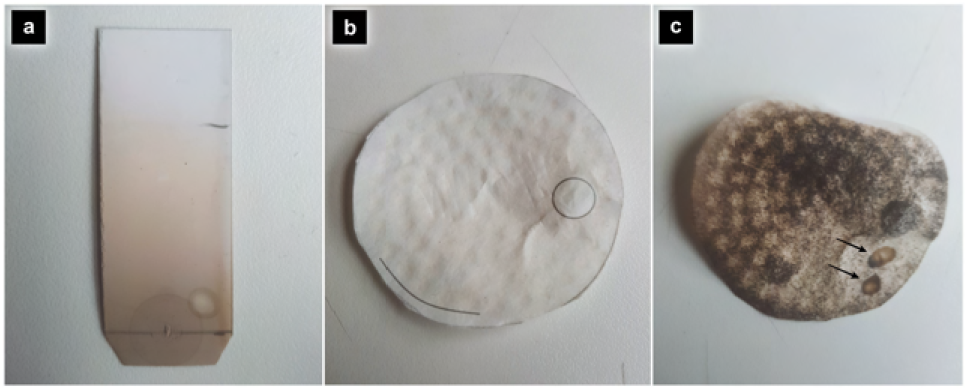
Solid-phase separation strategies for isolating FP-OTs. After Buchner solid extraction, the liquid was eluted on a TLC silica plate and sprayed with a mixture of H_2_SO_4_:methanol (1:1) and heated, but no sugar blotting was observed (a). In the cellulose filter, a circular white wax-like texture was found, believed to be the FP-OTs layered on top of the filter and unobtainable (b). The filter clearly interferes with the revealing solution (charred color), but some brownish spots characteristic of sugars can be observed (c).

#### Ideal solvent screening

The solubility of both native FucoPol and FP-OTs (*1a*) fibers was tested in different solvents for ^1^H-NMR preparation purposes. Solubility results are shown in Figure 6 and the corresponding weighed masses for ^1^H-NMR analysis are documented in Table 1. FucoPol in THF (Figure 6a) appears insoluble; fibers expanded, separated, and lost their dense and compact appearance previously observed in Figure 3e. This effect was also observed with 0.5 volumes of ethanol and isopropanol, but evaporation was more difficult. When dried (Figure 6b), its dense and compact fiber structure was restored, thus THF does not appear to chemically influence the polymer and can be used as a precipitating solvent. On the other hand, FP-OTs (*1a*) was tested in acetonitrile (Figure 6c), methanol (Figure 6d), chloroform (Figure 6e) and deuterated water (D_2_O, Figure 6f), showing high solubility in polar solvents, with D_2_O being the best fit, which implies potential compatibility with biological media.

**Table 1:** Weighed masses in solubility tests as solvent screening for ^1^H-NMR analysis.

| Sample | Sample mass | Solvent | Soluble |
| --- | --- | --- | --- |
| FP-OTs (*1a*) | 7.1 mg | Methanol | Partly |
|  | 9.6 mg | Chloroform | No |
|  | 9.3 mg | Acetonitrile | Yes |
| | 15.0 mg | $\text{D}_2\text{O}$ | Yes |

**Figure 6:**
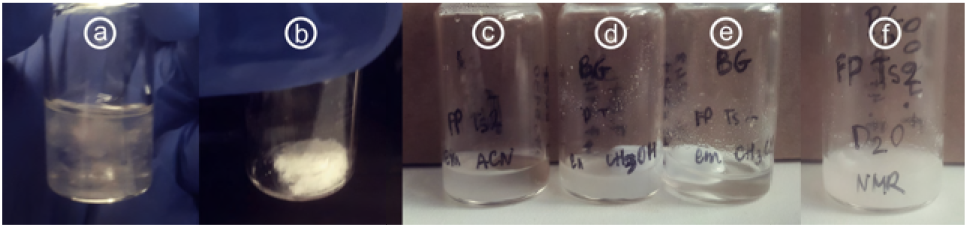
Solubility of native and tosylated FucoPol in different organic solvents. (a) FucoPol in THF. (b) FucoPol in THF (dried). (c) FP-OTs (*1a*) in acetonitrile. (d) FP-OTs (*1a*) in methanol. (e) FP-OTs (*1a*) in chloroform. (f) FP-OTs (*1a*) in D_2_O.

#### *ATR*-FTIR

Functional group analysis was carried out on all compounds to check for successful tosylation (Figure 7). The relevant vibrational changes of interest include double S=O bonds in the 1410– 1150 cm^−1^ range, aromatic C–C skeletal vibrations between 1625–1475 cm^−1^, and aromatic C–H near 3080–3030 cm^−1^. The most distinctive feature was a peak shift from 1602.3 cm^−1^, characteristic of native FucoPol, to 1627.9 cm^−1^, which is present in all other synthesized compounds. Although this shift does not fall under sulphate bond vibrations, it only occurs in compounds resultant from tosylation. Vibrations at 2620, 2605, 2531, and 2498 cm^−1^ show possible TEA contamination, as they only occur in the *b* compounds, tainted with its characteristic yellow tonality. A 1410–1380 cm^−1^ vibrational band also appears more strongly in *1b* and *2b*, hinting at a sulfonyl chloride S=O vibration from excess TsCl or SO_2_–OH vibrations from contaminant p-toluenesulfonic acid. In summary, FP-OTs (*1a*, the desired product) is conclusively contaminated with *1b* and *2b*. However, the 1602 to 1627 cm^−1^ shift may be indicative of successful tosylation or the presence of p-toluenesulfonic acid and triethylammonium p-toluenesulfate byproducts.

**Figure 7:**
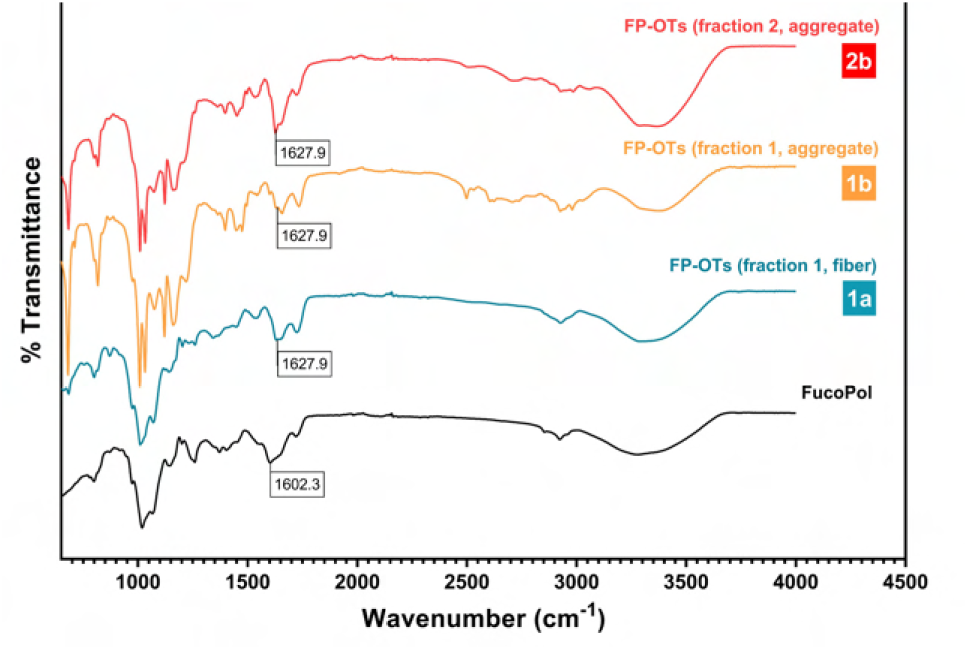
ATR-FTIR spectral comparison of FucoPol and ubiquitous tosylation products. Native FucoPol (black), compound *1a* (blue), compound *1b* (orange), and compound *2b* (red) are shown. A visible shift from 1602 to 1627 cm^−1^ is clearly observed in all tosylated derivatives.

#### ^1^H-NMR

To differentially diagnose plausible FucoPol tosylation, ^1^H-NMR analysis was performed with a focus on tosyl signals (Figure 8), showing evident differences at 7–8 ppm. Figures SI.1–SI.3 show the individual spectra and corresponding integrations of the relevant signals. For signal differentiation, the theoretical ^1^H-NMR spectra of the substances of interest—sugar monomer with free primary alcohol, tosylated sugar monomer, p-toluenesulfonic acid, and TEA—are shown in Figure SI.4. For the expected FP-OTs structure shown in Figure 2, see Figure SI.5. Compound *1a* (Figure 8, blue) shows four relevant signals: *δ* 7.92 (brs, 1H), 7.70 (s, 2H), 7.53 (brs, 1H), 7.38 (s, 2H). Considering the symmetry of the tosyl aromatic ring and the chemical microenvironment of its protons, only two doublet signals are expected near this region, instead of four. The two additional peaks suggest excess p-toluenesulfonic acid. Compound *2b* (Figure 8, red), present in fraction *2*, shows only two signals: *δ* 7.70 (d, J = 7.8 Hz, 1H) and 7.38 (d, J = 7.9 Hz, 1H). Considering that fraction *2* is a contaminant fraction which does not contain any tosylated polymer but excess reagent, due to its visual appearance, poor solubility, previous FTIR data, and these two signals, the differential inference is that the ^1^H-NMR signal pair corresponding to a sugar monomer tosylated in its primary alcohol position is the one with chemical shifts at 7.92 and 7.53 ppm, not found in *2b*. Their relative intensities also corroborate that excess p-toluenesulfonic acid is present, and from the crude NMR mass, FP-OTs amounts to 33%, similar to the 31.2% yield obtained from molar calculations.

**Figure 8:**
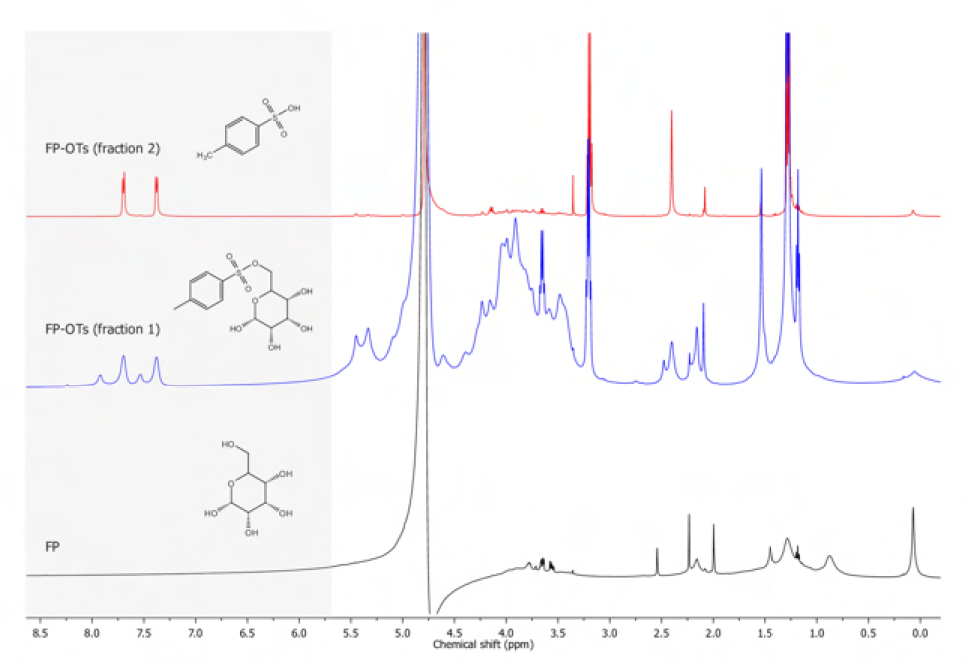
^1^H-NMR (D_2_O) of native FucoPol and synthesis derivatives. Native FucoPol (black), fraction *1* of FP-OTs (*1a* + *1b*, blue), and fraction *2* of FP-OTs (*2b*, red) are shown. The 7–8 ppm region is of particular interest for tosylation changes.

### FP-OTs synthesis pathway 2: COOH-targeted tosylation

The ubiquitous tosylation (pathway 1) and subsequent analysis confirmed the successful synthesis of the FP-OTs derivative, albeit of low yield (31.2%) due to existing byproducts. Thus, a second iteration of the derivatization focused on a more controlled synthetic approach to reduce excess reactants, and purification through dialysis to implement greener downstream processing and increase yield. Dialysis is a viable purification strategy as it leverages the high molecular weight of FucoPol (10^6^ Da range) to retain the FP-OTs inside the dialysis membrane and extract all other small-molecule organic contaminants in a one-step approach. The COOH-targeted synthesis procedure (Figure 2, pathway 2) also used a 0.25 wt.% solution of FucoPol, and the reaction mixture contained equimolar proportions of TsCl and TEA relative to the carboxyl contents of FP_*SRU*_ (1:1:1 TsCl:TEA:FP_*SRU*_), to target the single carboxyl group in the C_6_ of GlcA, located in the trimer branch of the polysaccharide. As before, the transparent beige appearance of native FucoPol converted to opaque white upon addition of TsCl and TEA. However, this synthetic approach did not produce the yellowish tone characteristic of excess TEA, and the visual appearance remained unchanged during the full extent of reactional stirring. Given the limiting reagent conditions used for pathway 2, two reaction times were tested to allow tosylation to occur in variable time windows: 18 h (overnight) and 96 h (4 days). In both cases, both reaction mixtures equilibrated as a two-phase separation post-synthesis, still revealing unreacted TEA (*ρ* = 0.726 g/mL) on top, the polymeric aqueous solution in the bottom (*ρ* ≈ 1 g/mL), and a cloudy, cotton-like interface (Figure 9) which might be either (i) a fraction of FP-OTs interacting with TEA due to acquired hydrophobicity, or (ii) an aqueous fraction supersaturated with triethylammonium salt.

**Figure 9:**
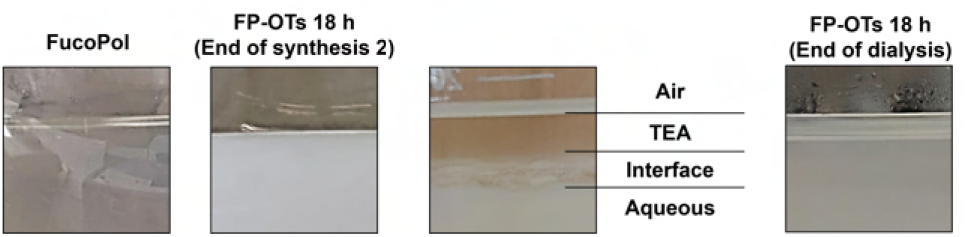
Temporal evolution of the appearance of a 0.25 wt.% FucoPol solution during COOH-targeted tosylation (pathway 2).

#### Dialysis monitoring

Both products of COOH-targeted tosylation, FP-OTs^18^ and FP-OTs^96^, were purified by dialysis using distilled water as the solvent exchange system. The temporal evolution of pH and Ω as control parameters for monitoring the extent of dialysis of tosylated derivatives is shown in Figure 10. The reaction mixture at the end of synthesis (Figure 10a, leftmost black datapoint) is highly alkaline due to the byproducts of tosylation, presenting pH 13.1 and Ω = 41.2 *µ*S/cm. The iterative dialysis procedure concomitantly reduces solution pH and Ω in a sigmoidal fashion, with an enhanced decrease in both properties in early dialysis steps and an attenuated purification in subsequent steps, which is expected from techniques relying on passive osmotic diffusion. After four solvent exchanges, aqueous FP-OTs^18^ is pH 9.4 and Ω = 0.5 *µ*S/cm, demonstrating conductivity levels similar to distilled H_2_O, the solvent exchange medium, indicating that dialysis is complete (Figure 10a, blue). However, the pH level is still alkaline and outside physiologically acceptable levels. Four additional solvent exchanges were carried out, although a fifth exchange was sufficient to lower pH to a biocompatible regime (Figure 10a, green). After eight dialysis steps, aqueous FP-OTs^18^ was pH 7.4 and Ω = 0.05 *µ*S/cm and freeze-dried. The dialysis of FP-OTs^96^ followed the same trend. After freeze-drying, the final yield of FP-OTs^18^ and FP-OTs^96^ was 62.0% and 64.8%, respectively. An average 63.4% yield was obtained with dialysis, double that of solvent washing techniques (31.7%), revealing a superior purification method with no byproduct contaminants.

**Figure 10:**
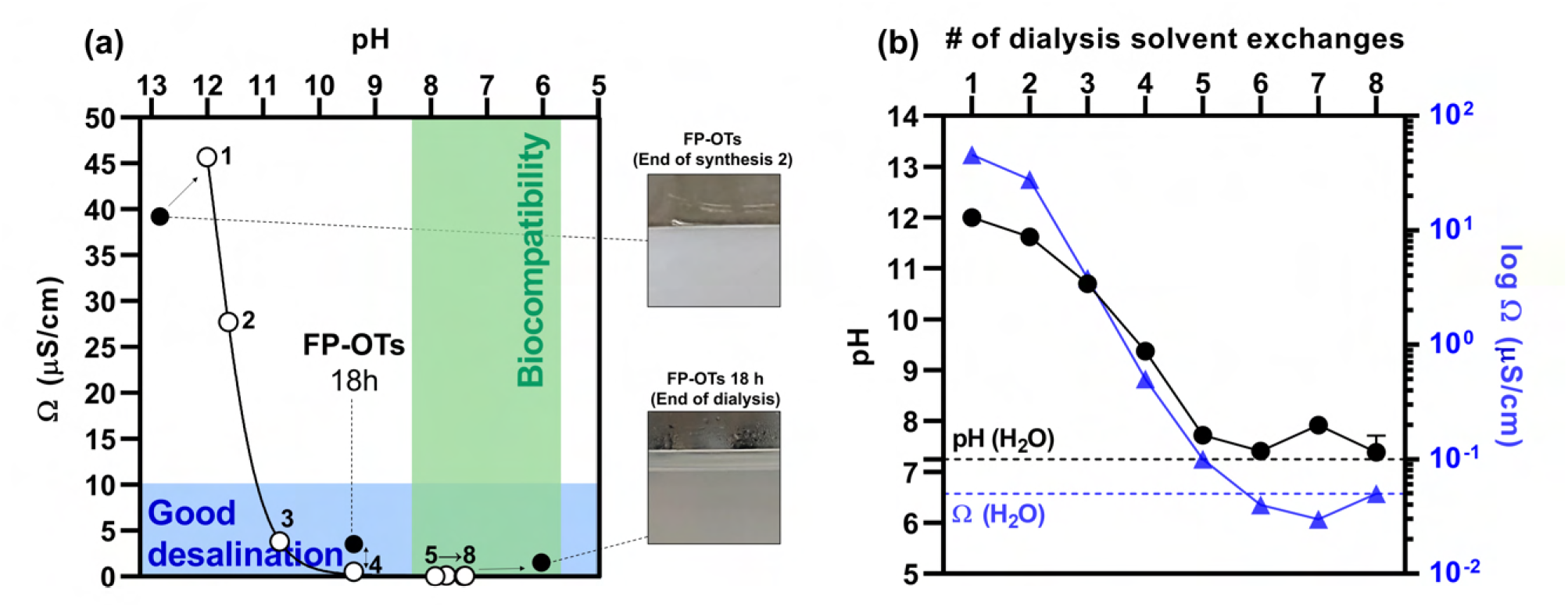
Dialysis monitoring of the purification of FP-OTs^18^, obtained from COOH-targeted tosylation. (a) pH–Ω matrix representation of the evolution of dialysis over a maximum of eight solvent exchanges. Black datapoints, from left to right, represent reaction mixtures at the end of synthesis, the final FP-OTs^18^ product used for further analysis (four solvent exchanges), and a supplementary FP-OTs^18^ fraction representing the end of dialysis (eight solvent exchanges). White datapoints represent the pH and Ω of the aqueous FP-OTs^18^ inside the dialysis membranes at each solvent exchange checkpoint. (b) Complementary pH and Ω data for the FP-OTs^18^ aqueous product inside the dialysis membranes, with distilled H_2_O levels used as reference.

#### Physicochemical properties and molecular weight

The physicochemical properties and *M*_w_ of both FP-OTs variants re-dissolved in *d*H_2_O, post-freeze-drying, are shown in Table 2. A significant reduction in solution pH and Ω was observed for tosylated FucoPol derivatives compared to native FucoPol, which is suggestive of successful tosylation due to increased hydrophobicity. The similar properties of FP-OTs^18^ and FP-OTs^96^ also indicate that 18 h of reaction time is sufficient for tosyl substitution to be carried out. The *M*_w_ of FP-OTs derivatives revealed no significant change in structure length. FP-OTs^18^ and FP-OTs^96^ had *M*_w_ values of 1.14*±*0.002 MDa (*p* = 0.845) and 1.10*±*0.046 MDa (*p* = 0.692), respectively, presenting no statistically significant difference from native FucoPol (1.25*±*0.005 MDa).

**Table 2:** Physicochemical properties of native FucoPol, COOH-targeted FP-OTs variants after freeze-drying and redissolution, and the approximate average of a consortium of bio-based polysaccharides used in biological cryopreservation studies [7], for reference.

| Sample | pH | Ω (μS/cm) | $M_w$ ($\times 10^6$ Da) | $M_n$ ($\times 10^6$ Da) | PDI |
| --- | --- | --- | --- | --- | --- |
| FP | 6.73 ± 0.03 | 798.9 ± 130.6 | 1.25 ± 0.005 | 0.92 ± 0.03 | 1.35 ± 0.04 |
| FP-OTs<sup>18</sup> | 4.76 ± 0.02 | 331.4 ± 6.1 | 1.14 ± 0.002 | 0.74 ± 0.02 | 1.55 ± 0.03 |
| FP-OTs<sup>96</sup> | 5.32 ± 0.09 | 315.4 ± 30.0 | 1.10 ± 0.046 | 0.69 ± 0.23 | 1.67 ± 0.34 |
| Other polys. | 5.77 ± 0.34 | 647.7 ± 37.9 | – | – | – |

#### Rheology of tosylated derivatives

The unaffected molecular weight hinted at no degradation as a consequence of synthesis. Thus, the rheological behavior of native and tosylated FucoPol structures was probed to assess their structural integrity (Figure 11). The characteristic shear-thinning behavior of native FucoPol observed in Figure 11a was partially preserved in the tosylated derivatives, but an extensive loss of viscosity was observed. At rest, FP-OTs^18^ and FP-OTs^96^ expressed, respectively, only 40.6% and 31.1% of the zero-shear viscosity (*η*_0_) of native FucoPol. This indicates that although no significant *M*_w_ decrease occurred, the internal structuration of tosylated FucoPol derivatives was affected. This is consistent with a gradual decrease in *M*_n_ and concomitant increase in polydispersity index (PDI) with reaction time, revealing some breakage of interchain bonding. The insertion of a tosyl group in the C6 carboxyl region may lead to steric hindrance and hinder intermolecular chain arrangement and non-covalent electrostatic interactions that confer robustness to the structure, and therefore, the rheological properties natively observed. Contrary to native FucoPol, neither FP-OTs variant achieved a gel state, due to the inexistence of a *G*′–*G*″ curve crossover and the dominance of the loss modulus *G*″. An unchanged molecular weight post-synthesis, accompanied by a loss of zero-shear viscosity and the inability to produce a gel state are related to macromolecular re-structuring rather than chemical degradation, as express consequences of a trade-off between inserting a hydrophobic group of high steric hindrance (tosyl) and removing a hydrophilic group of low steric hindrance (carboxyl). The loss of overall electronegativity reduces chemical affinity with solvent water but also precludes the generation of a strong network of interchain non-covalent interactions contributive to a gel state.

**Figure 11:**
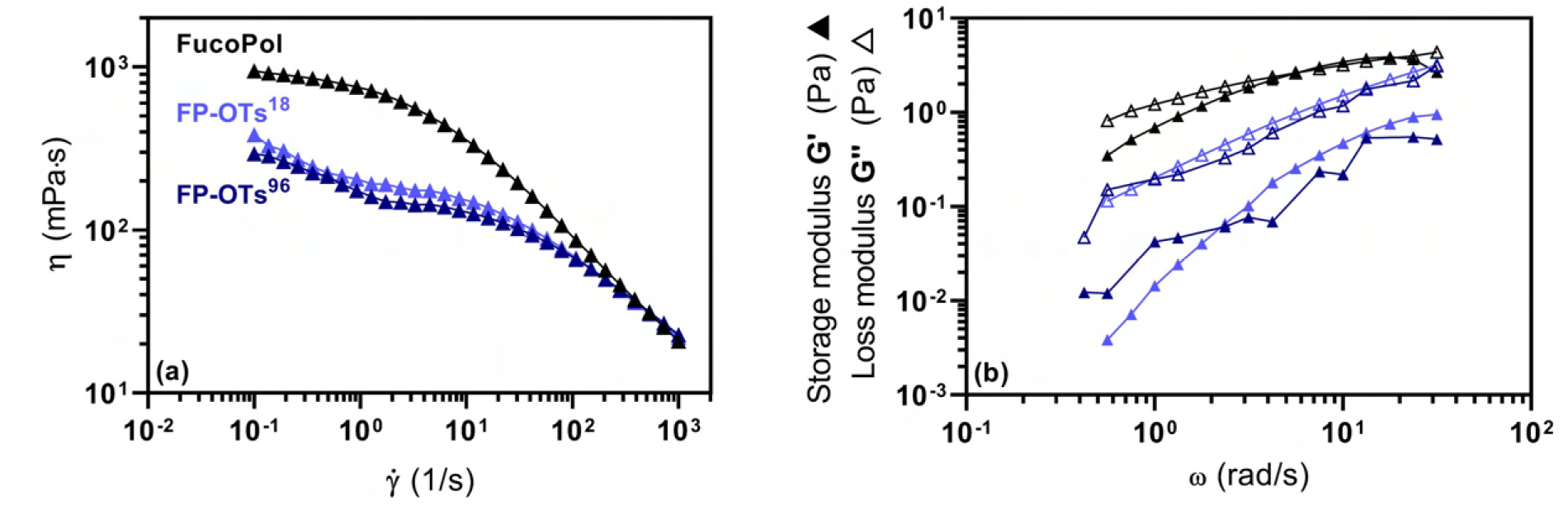
Rheological behavior of native and tosylated FucoPol structures. (a) Flow curves of aqueous solutions of 0.5 wt.% FucoPol (black), FP-OTs^18^ (light blue) and FP-OTs^96^ (dark blue), at 4 °C. (b) Frequency sweeps of the corresponding samples, at 4 °C, showing both *G*′ (storage) and *G*″ (loss) moduli.

#### Cytotoxicity

To assess the cryoprotective potential of the newly tosylated FucoPol structures, a preliminary in vitro assay to discard potential cytotoxicity as a cause for cellular death was carried out. Figure 12 shows the metabolic viability of Vero cells in the presence of FP-OTs^18^ and FP-OTs^96^ for 48 h at 37 °C. Despite some apparent cytotoxicity for FPOTs^18^ at 0.13 and 0.06 wt.%, both tosylated derivatives at their highest concentration tested (0.25 wt.%) are within the healthy viability range. Thus, the tosylation of FucoPol was considered to be a biologically safe methodology for polysaccharide functionalization.

**Figure 12:**
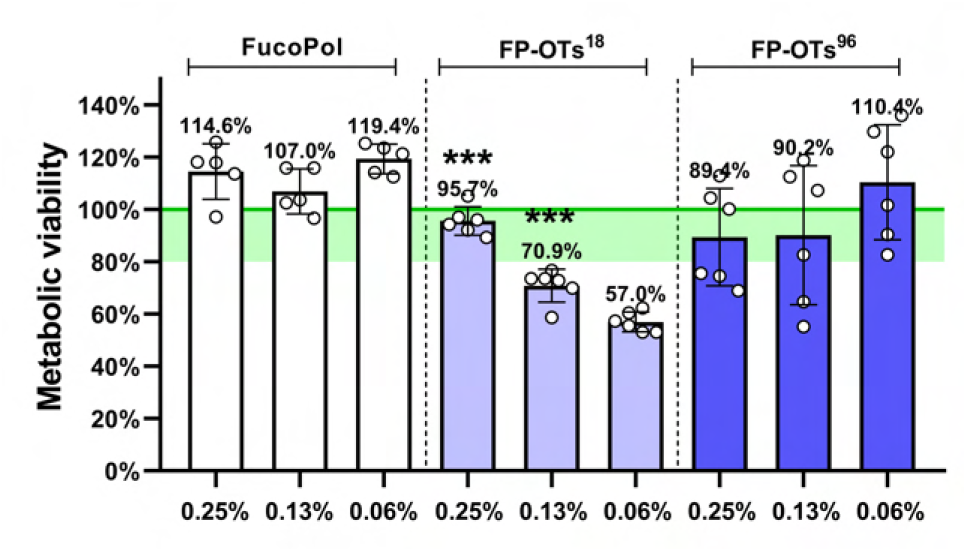
Vero cell metabolic viability cytotoxicity assay of FP-OTs derivatives. Statistical significance was determined using a Brown-Forsythe and Welch ANOVA test, by comparing each corresponding concentration between FPOTs^18^ or FP-OTs^96^ data and the native FucoPol benchmark. The green shaded region represents the interval (80–100%) for which a substance exposed to the seeded cells is, by convention, deemed biocompatible.

#### Cryoprotective activity

The synthesis of tosylated FucoPol derivatives in their purified form intended to assert the significance of knocking out UAs from a polysaccharide structure to measure a change in cryoprotective activity. Figure 13 shows the cryoprotective performance of purified, biocompatible FP-OTs^18^ and FP-OTs^96^ dissolved in DMEM culture medium, co-supplemented with 10% DMSO. All tosylated FucoPol derivatives, regardless of concentration or reaction time, effectively lost the cryoprotective activity inherent to their native counterpart. Both FP-OTs^18^ and FP-OTs^96^ demonstrated suboptimal cell survival, even in the presence of 10% DMSO, and below the threshold for standalone 10% DMSO, revealing the lack of any cryoprotective activity. Compared to the 0.7, 1.5 and 2.9-fold change in Vero cell metabolic viability experienced in the presence of native FucoPol, FP-OTs^18^ resulted in 0.3, 0.7 and 0.1-fold changes and FP-OTs^96^ resulted in 0.2, 0.3 and 0.3-fold changes, by increasing order of concentration.

**Figure 13:**
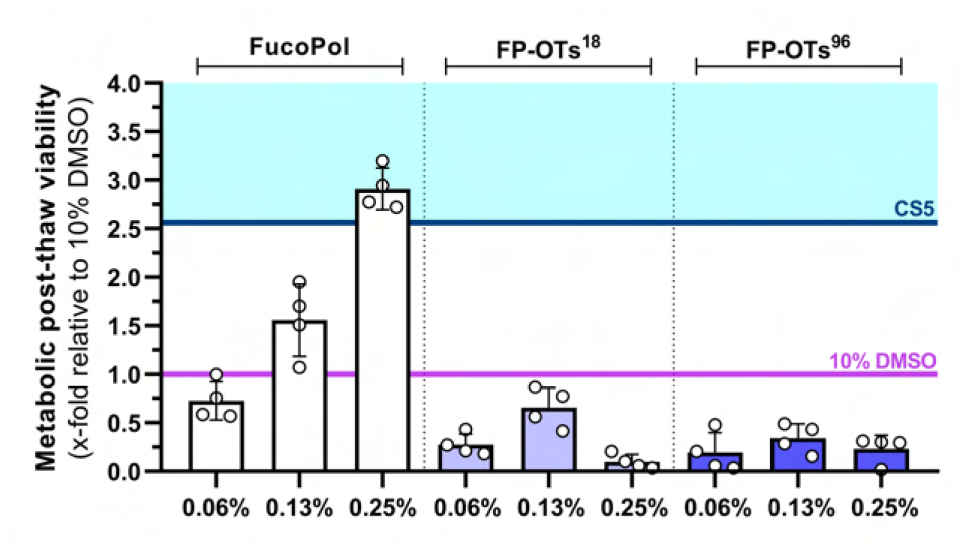
Cryoprotective performance of native and tosylated FucoPol derivatives, FP-OTs^18^ and FP-OTs^96^. The post-thaw metabolic viability of Vero cells is shown. The blue and purple thresholds correspond to the cryoprotective performance of CryoStor^TM^ CS5 and standalone 10% DMSO, respectively, as experimental controls.

## Discussion

The tosylation of the bio-based cryoprotective polysaccharide FucoPol was explored for two main purposes. First, to interrogate the validity of the main paradigm in chemical cryobiology, in which cryoprotective activity arises from an acquired permanent dipole moment, such as uronic acid moieties in polysaccharides. Second, the functionalization of polysaccharides has scarce literature because their complexity often hinders synthesis and a complete characterization is strenuous, but the ability to enhance cryoprotective properties by additive synthesis generates significant interest. Here, the chosen modification of native FucoPol structures was tosylation. The tosyl moiety is an excellent leaving group, thus allowing for its easy displacement by SN_2_ reactivity while using tosylated structures as a building block for other products. The O-tosylation mechanism is a nucleophilic attack by the TsCl to a *sp*_2_ carbon, which displaces the –OH group and forms HCl and p-toluenesulfonic acid as byproduct, thus the need for TEA as counteracting base to acid formation, to avoid acid hydrolysis of the polymer chains. The initial 10 min of reaction were performed at 4 °C to decrease the nucleophilic reactivity of water and allow O-tosylation to occur without competition. Due to steric effects, tosylation is expected to occur only in the primary alcohols of the sugar backbone, particularly on the C_6_-linked carboxyl of *β*-D-GlcA*p* in the trimer branch (Figure 2). This work focused on the proof-of-concept synthesis, downstream optimization and reassessment of physical, chemical and biological properties. The FP-OTs derivative was successfully synthesized by two pathways: by ubiquitous tosylation using a 1:7:7 FucoPol:TsCl:TEA excess; and by COOH-targeted tosylation, which used a 1:1:1 FucoPol:TsCl:TEA ratio stoichiometrically balanced with the single carboxyl moiety in the structural repeating unit of FucoPol. Signs of successful tosylation included an ATR-FTIR signal shift from 1602.3 to 1627.9 cm^−1^, two broad singlet at *δ* 7.92 and 7.53 ppm in ^1^H-NMR spectra, reduction in solution conductivity from ca. 800 to 300 *µ*S/cm and a 30–40% reduction in zero-shear viscosity from loss of intermolecular chain non-covalent interactions due to enhanced steric hindrance. Dialysis proved to be a better purification method for FP-OTs derivatives, yielding an average 63.4% mass dry weight compared to the average 31.7% yield using organic solvent evaporation and washing techniques, which nevertheless would accrue contaminant byproducts in the freeze-dried product. All aforementioned identifiers of successful tosylation relate to a product that is more hydrophobic, less anionic and structurally less robust at a macromolecular scale, which is suggestive of a decreased cryoprotective activity. Although all FP-OTs derivatives were biocompatible, a drastic loss of cryoprotective activity was observed, leading to an antagonistic effect in cell survival. Freezing media containing FP-OTs generated PTV fold-changes between 0.1–0.7, a lower performance than that of standalone 10% DMSO. These results confirmed that uronic acids are a key player in polysaccharide-based cryopreservation, thus supporting the dogma of electronegativity as a source of ice growth disruption and previous research [7]; but also revealed that it was the subsequent structural modifications arising from a removal of the carboxyl group that led to the total loss of cryoprotective activity, as the inability of generating a gel state at 4 °C like native FucoPol, as also shown before [11], stands as the most probable cause for the lack of efficient cryoinjury avoidance. Although UAs were pinpointed as an indispensable moiety in cryoprotective polysaccharides, the FP-OTs derivative can be rationalized as an intermediate building block for further cryoprotective-enhancing modifications or fluorescence strategies. The research flexibility that polysaccharide tosylation brings provides exciting opportunities to explore a library of promising small molecules, and acts as a stepping-stone in bottomup quantitative structure-activity relationship analysis with an organic chemistry approach. The latest attempt at combining organic chemistry, carbohydrate chemistry and cryopreservation involved the dephosphorylation of a mannan [19], a subtractive strategy. In what concerns addition, the anchoring of sulphate-based moieties mimicking DMSO to a polymeric backbone proved unsuccessful: sulfide anchors were cytotoxic to cells, and sulfoxide anchors did not show any cryoprotective effect [22]. Amino acid substituents are particularly interesting due to their unveiled properties in crystal interaction. *Colwellia psychrerythraea* 34H is a psychrophilic bacterium that produces a seminal EPS with promising cryoprotective properties due to its unique amino acid substituents in the structure [23], which draw some parallelisms to the known effects of antifreeze proteins and their structure-function relationships to ice binding [24]. Nitrogen chemistry using azide linkers to anchor other substituents has also been implemented in photoisomerization strategies, to create photoactive molecular switches that can hinder or allow ice recrystallization to occur [25]. Using fluorophores as substituents would provide an exciting opportunity to probe polysaccharides in the cellular milieu, during all stages of the freeze-thaw process, *e*.*g*., to monitor the spatial arrangements of polysaccharide molecules in-between the crystal lattice and the cell surface, or to assess the growth morphology of boundary-constrained polymorphs in the presence of a polysaccharide.

## Conclusion

The synthesis of UA knock-out tosylated derivatives from the cryoprotective polysaccharide FucoPol envisioned challenging the dogma of electronegativity as the sole predictor of ice growth disruption. The SN_2_ tosylation of the single axial carboxyl (63.4% yield) followed by green dialysis is a quick and easy derivatization able to generate a polysaccharide variant that is more hydrophobic, less anionic and structurally less robust at a macromolecular scale. Although biocompatible, the loss of polyanionicity which cascaded into the inability of maintaining rheological integrity and forming a gel-state near hypothermia resulted in a complete loss of cryoprotective activity. These results emphasize the critical importance of UAs in polysaccharide structures to elicit a biological cryoprotection benefit not simply for their standalone electronegativity, but also for their participation in inter-chain interactions which sustain the conformational integrity necessary for bioactivity. The continued advancement of new polysaccharide functionalization strategies, although strenuous from a synthetic and analytical perspective, is indispensable because the functional enhancement of current cryoprotectants by building-block additive synthesis generates significant interest.

## CRediT Authorship

B.M.G. conceptualized study, performed experiments, data analysis, wrote manuscript. F.F., J.L., J.S. reviewed manuscript, supervised, funded. All authors have read and agreed to the published version of the manuscript.

## Data Availability Statement

The data that support the findings of this study are available from the corresponding author upon request.

