## Supplementary Information for "Tosylation-mediated uronic acid knockout from cryoprotective FucoPol revealed the importance of polyanionicity in ice growth disruption"

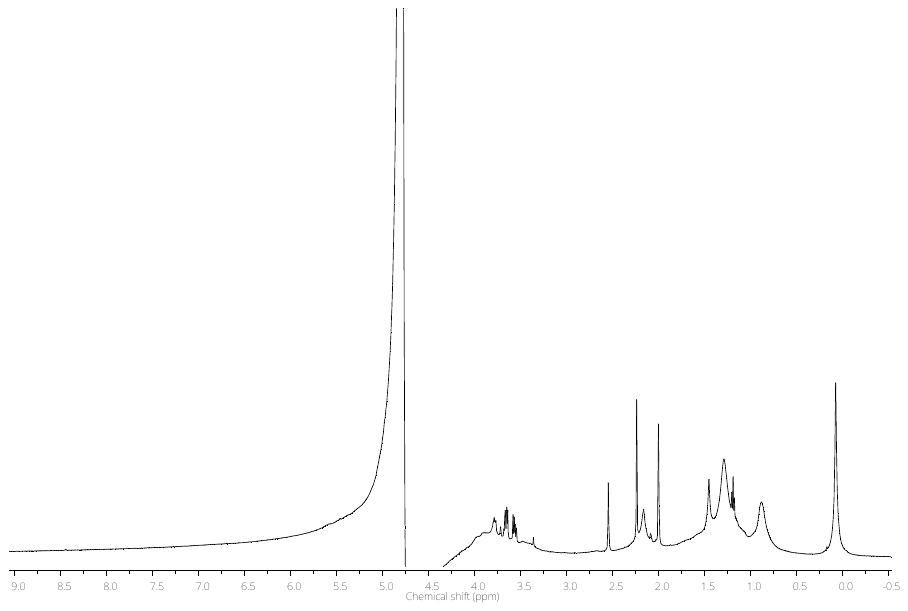

Figure SI.1 – ^1^H-NMR (D_2_O) spectrum of native FucoPol.

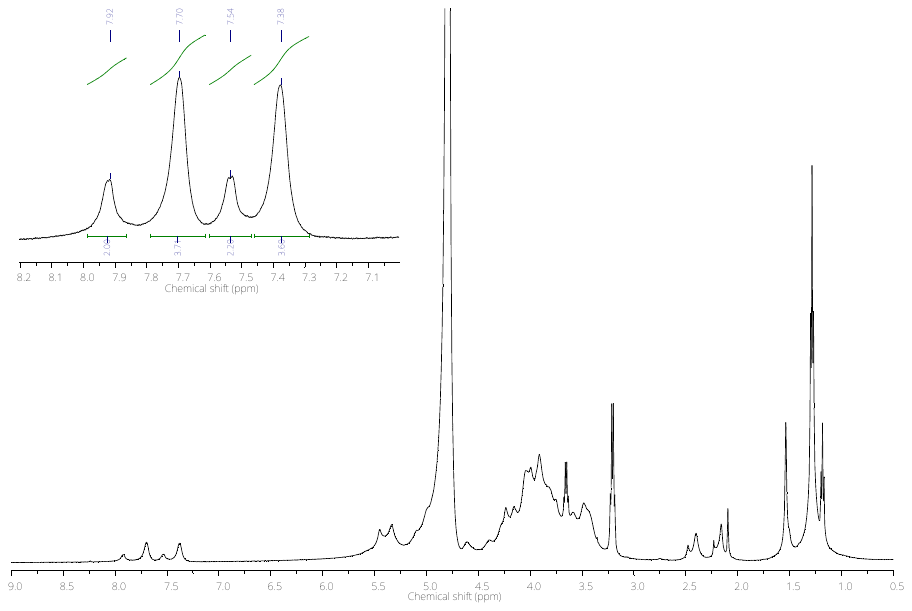

Figure SI.2 – ^1^H-NMR (D_2_O) spectrum of fraction 1 of FP-OTs (1a + 1b). In inset, the 7–8 ppm region related to tosyl aromatic rings.

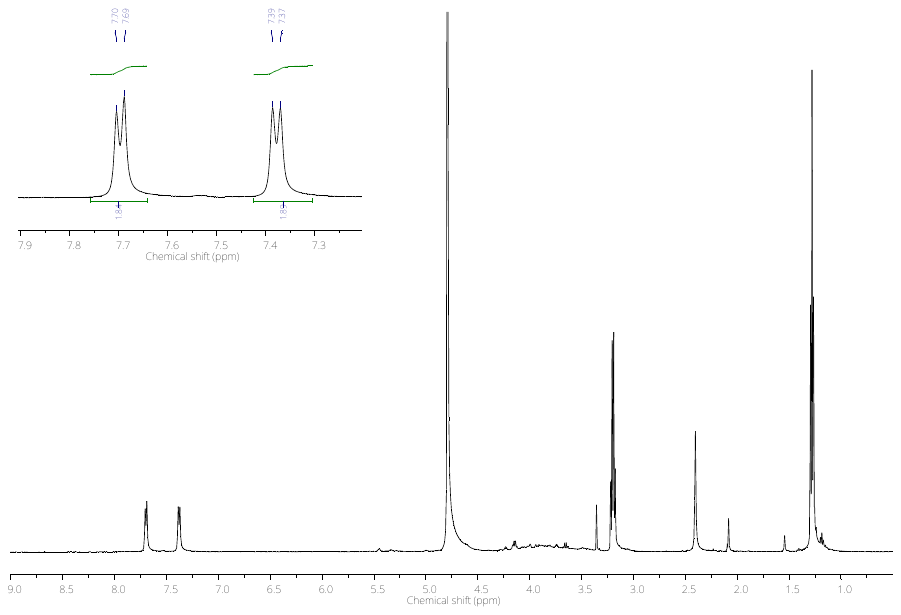

Figure SI.3 – ^1^H-NMR (D_2_O) spectrum of fraction 2 of FP-OTs (2b). In inset, the 7–8 ppm region related to tosyl aromatic rings.

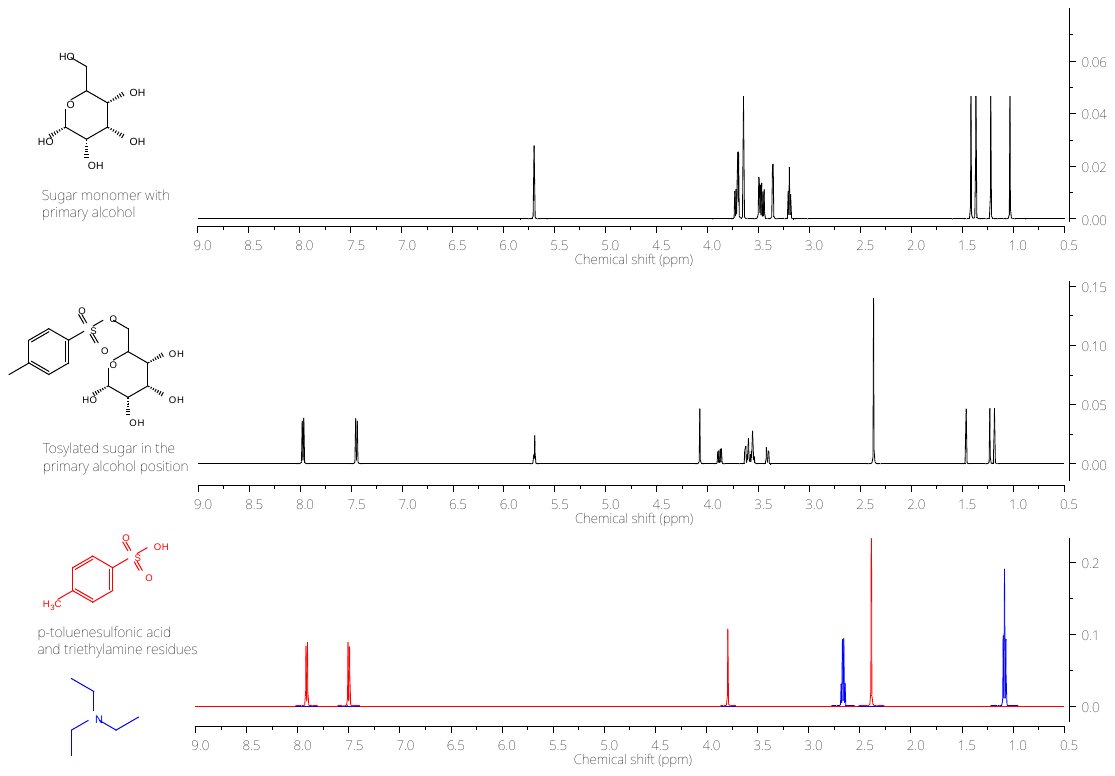

Figure SI.4 – Theoretical ^1^H-NMR (CHCl_3_) spectra of all chemical species involved in FucoPol tosylation. From top to bottom, the core sugar monomer with a free primary alcohol, tosylated sugar monomer, p-toluenesulfonic acid (red) and triethylamine (blue). The simulation was performed in MestReNova 6.0.2.

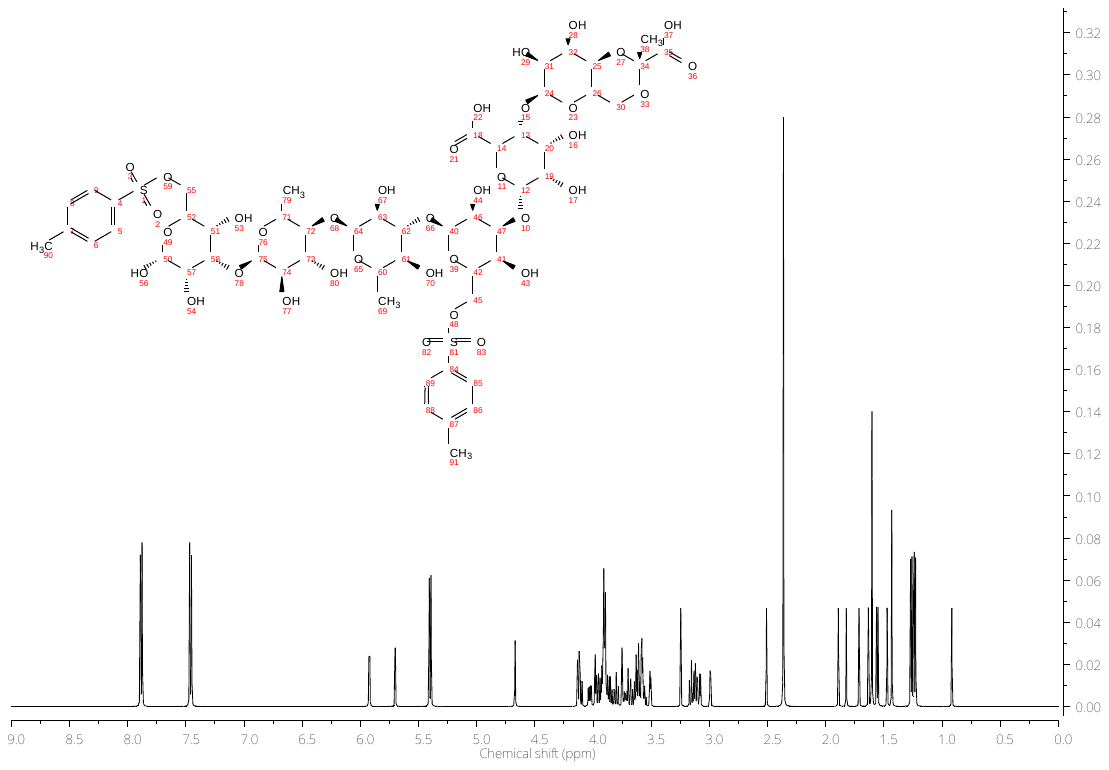

Figure SI.5 – Theoretical ^1^H-NMR (CHCl_3_) spectra of the FP-OTs structural repeating unit. The simulation was performed in MestReNova 6.0.2.
